# Harnessing Diacylglycerol-Terminated Cationic Oligomers for Next-Generation Antibacterial Therapeutics

**DOI:** 10.64898/2026.04.01.715743

**Authors:** Qiaomingxuan Liu, Shiwei Zhang, Margaret Pywell, Alysha G. Elliott, Holly Floyd, Johannes Zuegg, Jessica R. Tait, John F. Quinn, Michael R. Whittaker, Muhammad Bilal Hassan Mahboob, Cornelia B. Landersdorfer

## Abstract

Cationic polymers, which mimic the structure of antimicrobial peptides (AMPs), are increasingly recognized as promising antimicrobial materials. Here, we report the synthesis and evaluation of a new class of cationic lipid-terminated oligomers (CLOs), comprised of 2C_18_-hydrophobic lipid tails, and short oligomeric cationic chains synthesised via Cu(0)-mediated reversible-deactivation radical polymerization (RDRP). Two 2-vinyl-4,4-dimethyl-5-oxazolone (VDM) oligomers with degrees of polymerization (DP) of 20 or 50 were synthesized using the lipid functional initiator (R)-3-((2-bromo-2-methylpropanoyl) oxy)propane-1,2-diyl dioctadecanoate (2C_18_-Br). Post-polymerization modification of the pendant oxazolone moieties was carried out using reactive amines, including *N*-Boc-ethylenediamine (BEDA) and *N,N*-dimethylethylenediamine (DMEN). Subsequent deprotection of the BEDA groups and quaternization of DMEN groups enabled the synthesis of six functional CLOs exhibiting distinct cationic functionalities. Antimicrobial assays against a panel of WHO bacterial and fungal priority pathogens (methicillin-resistant *Staphylococcus aureus* [MRSA], *Escherichia coli, Klebsiella pneumoniae, Acinetobacter baumannii, Pseudomonas aeruginosa, Candida albicans,* and *Cryptococcus neoformans*) revealed that these CLOs exhibited potent and selective structure-dependent antibacterial activity, particularly against MRSA, with minimum inhibitory concentrations (MICs) in the clinically relevant range, below 4 µg mL^-1^, comparable to antibiotics vancomycin and colistin. Among these, BEDA-functionalized CLOs demonstrated the strongest antimicrobial profile, which was significantly increased by increasing DP, as evidenced by a reduction in MIC values from 64 µg mL^-1^ (for DP20) to ≤ 4 µg mL^-1^ (for DP50) against *A. baumannii*. Biocompatibility assays against red blood cells and HEK293 cells indicated negligible toxicity, with haemolytic (HC_50_) and cytotoxic (CC_50_) values exceeding 512 µg mL-1 across all CLOs. All CLOs displayed minimal activity against *C. albicans* (MIC ≥ 512 µg mL^-1^). In contrast, activity against *C. neoformans* was influenced by both cationic functionality and DP, with DMEN-based CLOs exhibited superior antifungal activity at higher DP relative to their BEDA-based counterparts. Most CLOs displayed high selectivity (SI) toward MRSA (SI >128), while 2C_18_-O(BEDA)_50_ exhibited the broadest spectrum, showing potent antimicrobial activity and high selectivity against *E. coli* (MIC ≤ 4 µg mL^-1^, SI ≥ 128), *A. baumannii* (MIC ≤ 4 µg mL^-1^, SI ≥ 128), and MRSA (MIC ≤ 4 µg mL^-1^, SI ≥ 128), along with moderate activity against *P. aeruginosa* (MIC = 32 µg mL^-1^, SI > 16). Taken together, these findings elucidate the combined influence of end-group lipidation, cationic functionality, and polymer length in modulating antimicrobial activity, thereby establishing 2C_18_-terminated CLOs as a rationally tunable and biocompatible platform for antimicrobial material development.

## Introduction

Antimicrobial resistance (AMR) is escalating worldwide and is now regarded as one of the most serious threats to human and animal health [1]. In 2019, bacterial AMR was associated with more than 4.9 million deaths, including 1.27 million directly attributable fatalities, and projections estimate this number could rise to 10 million annually by 2050 without effective intervention [2, 3]. Central to this crisis are the ESKAPE pathogens (*Enterococcus faecium, Staphylococcus aureus, Klebsiella pneumoniae, Acinetobacter baumannii, Pseudomonas aeruginosa,* and *Enterobacter spp.*), which account for the majority of hospital-acquired infections and AMR-related mortality [4]. These organisms were first recognised as critical threats by the Infectious Diseases Society of America (IDSA) in 2009 [5, 6]. This classification has since been reinforced by the World Health Organization (WHO) through its updated priority pathogen list in2024, and by the U.S. Centers for Disease Control and Prevention (CDC) in 2019 [7, 8]. Collectively, these frameworks guide research prioritisation, inform antimicrobial stewardship strategies, strengthen surveillance systems, and accelerate the development of novel therapeutics [9, 10].

Within this group, methicillin-resistant *Staphylococcus aureus* (MRSA) and *A. baumannii* exemplify the growing AMR crisis [11, 12]. MRSA, a Gram-positive pathogen resistant to nearly all β-lactam antibiotics, continues to cause severe systemic infections with limited treatment options [13–15]. Meanwhile, *A. baumannii*, a Gram-negative bacterium that can form robust and persistent biofilms, is increasingly associated with multidrug resistance, treatment failure, and high mortality [16–20]. These pathogens highlight the declining effectiveness of conventional antibiotics, which remain central to modern medicine but are increasingly compromised by resistance, driven in part by misuse. [21].

Efforts to address antimicrobial resistance (AMR) have largely focused on modifying existing antibiotic scaffolds, often yielding derivatives that target pathways for which resistance mechanisms are already established[22, 23]. As a result, many candidates fail to progress through clinical development due to insufficient mechanistic novelty [24]. These limitations underscore the urgent need for innovative approaches that can circumvent established resistance mechanisms while maintaining efficacy and safety [25].

One promising strategy to tackle AMR is the use of antimicrobial polymers, particularly antimicrobial peptide (AMP) mimicking polymers [26–30]. Natural AMPs, as key mediators of innate immunity, exhibit potent antimicrobial activity; however, their clinical translation is hindered by poor biocompatibility, rapid *in-vivo* degradation, systemic toxicity, and high production costs [31–34]. Synthetic AMP-mimicking polymers, which co-opt the structural features of AMPs into synthetic polymers, provide an attractive alternative, offering structural tunability, enhanced stability, and cost-effective production [35–38]. Within this class, non-lipidated polymers act mainly through electrostatic disruption of microbial membranes, but their efficacy is often restricted by limited potency and cytotoxicity at elevated concentrations [39–43]. To overcome these shortcomings, lipidated cationic antimicrobial polymers, most notably cationic lipidated oligomers (CLOs), have been designed to integrate cationic charge with hydrophobic lipid moieties [44–48]. This combination facilitates simultaneous electrostatic binding and hydrophobic membrane insertion and destabilisation, thereby enhancing antimicrobial potency, stability and selectivity, while reducing toxicity [49, 50]. Moreover, CLOs exhibit structure-dependent broad-spectrum antimicrobial activity and strong compatibility with mammalian cell membranes, positioning them as a highly promising platform with advantages over non-lipidated counterparts for clinical translation.[37, 51, 52].

*Grace et al.* reported the use of azlactone-based chemistry to develop CLOs as a modular platform, enabling rapid and parallel functionalization with diverse nucleophiles under mild and atom-efficient conditions. This strategy allows facile incorporation of functional groups without generating by-products, greatly accelerating antimicrobial screening [53]. Studies have revealed that tuning the hydrophobicity of CLOs modulates their biological activity, while increased hydrophobicity enhances membrane penetration, excessive hydrophobicity induces haemolysis [54, 55]. Our previous work demonstrated that cholesterol and didodecanoate-based lipid tails conferred potent antifungal activity with acceptable toxicity, validating the role of lipid moieties in modulating CLO function [53, 56]. In this work, we focused on diacylglycerols (DAGs) such as 1,2-distearoyl-sn-glycerol (2C_18_), which contain two saturated C_18_ fatty acid chains. These long-chain saturated fatty acids readily insert into and destabilize bacterial membranes, thereby enhancing bactericidal activity [57, 58]. Thus, incorporating DAG motifs into CLOs enables systematic exploration of how lipid architecture influences their structure–activity relationships and antibacterial efficacy.

In this study, we report the synthesis and evaluation of AMP-mimicking CLOs incorporating a terminal 1,2-(C18:C18) DAG (2C_18_) group as a membrane-permeating agent. A well-defined library of VDM-based oligomers was synthesized via Cu(0)-mediated RDRP using 2C_18_-Br as the initiator, targeting degrees of polymerization (DP) of 20 and 50 **(Scheme 1)**. These oligomers were ring-opened to carry primary amine, tertiary amine, and quaternary ammonium functionalities **(Scheme 2).** This library was screened for antimicrobial activity, and minimum inhibitory concentrations (MICs) against WHO bacterial and fungal priority pathogens, including MRSA, *E. coli*, *K. pneumoniae*, *A. baumannii*, *P. aeruginosa*, *Candida albicans*, and *Cryptococcus neoformans*. Cytotoxicity to mammalian cells was further assessed using human red blood cells (RBCs) and HEK293 cells. Together, these evaluations provide critical insights into how structural variations in CLO-based antimicrobial polymers influence their antimicrobial efficacy, selectivity and cytocompatibility, thereby guiding their potential therapeutic applications.

## Experimental

### Materials

2-2-Vinyl-4,4-dimethyl-5-oxazolone (VDM) was synthesised as previously described [59], distilled under vacuum, and stored at -20 °C. Tris(2-(dimethylamino)ethyl)amine (Me_6_TREN) was synthesised according to the method of *Ciampolini et al* [60]. Acetone (Ajax FineChem), aluminium oxide (activated, neutral, Sigma-Aldrich), copper (II) bromide (CuBr_2_, Sigma-Aldrich, 99%), dichloromethane (DCM, Merck) *N,N*-dimethylacetamide (DMAc, Sigma-Aldrich, for HPLC, ≥99%), dimethylformamide (DMF, Merck Millipore), methanol (Ajax FineChem), *N*-Boc-ethylenediamine (BEDA, Sigma-Aldrich), *N,N*-dimethylethylenediamine (DMEN, Sigma-Aldrich), triethylamine (TEA, Sigma-Aldrich), trifluoroacetic acid (TFA, Sigma-Aldrich), iodomethane (Sigma-Aldrich), magnesium sulfate anhydrous (MgSO_4_, Sigma-Aldrich) were used as received. Copper wire was pre-activated by exposing it to sulfuric acid for 10 min, then rinsed with water to remove excess acid and stored in a closed bottle at room temperature. Dialysis was conducted using Cellu Sep H1 MWCO 2,000 Flat Width 38mm dialysis membranes (Membrane Filtration Products, Inc).

The bacterial and fungal strains used were methicillin-resistant *Staphylococcus aureus* (MRSA) ATCC 43300, *Escherichia coli* ATCC 25922, *Klebsiella pneumoniae* ATCC 700603, *Acinetobacter baumannii* ATCC 19606, *Pseudomonas aeruginosa* ATCC 27853, *Candida albicans* ATCC 90028, and *Cryptococcus neoformans* H99 ATCC 208821. The cell line used in the cytotoxicity assay was HEK293 human embryonic kidney cells, ATCC CRL 1573. Human whole blood was acquired from the Australian Red Cross Blood Service.

## Methods

### Synthesis of (R)-3-((2-bromo-2-methylpropanoyl)oxy)propane-1,2-diyl dioctadecanoate (2C_18_-Br) initiator

18:18-Diacylglycerol initiator (2C_18_-Br) was synthesised as previously reported [61]. Briefly, 1,2-*O*-dioctadecyl-sn-glycerol (1.31g, 2.19 mmol) was dissolved in 20 mL DCM in a 500 mL round-bottom flask and stirred for 15 mins at room temperature. Triethylamine (1.56 mL, 1.14 g, 11.2 mmol) was then added to the reaction mixture, and the solution was cooled in an ice bath before α-bromoisobutyryl bromide (0.54 mL, 1.0 g, 4.38 mmol) was added dropwise. The solution was left to stir at room temperature for 24 h. The resulting brown solution was then diluted with DCM and washed 3 times with water, followed by brine. The DCM layer was then dried with anhydrous MgSO_4_, filtered, and the solvent removed under reduced pressure. The crude oil was then purified via silica gel chromatography using DCM as the solvent. The purified initiator was isolated as a straw-coloured oil after drying in a vacuum oven overnight. ^1^H nuclear magnetic resonance (^1^H NMR) analysis confirmed the structure of 2C_18_-Br.

### (R)-3-((2-bromo-2-methylpropanoyl)oxy)propane-1,2-diyl dioctadecanoate (2C_18_-Br)

^1^H-NMR CDCl_3_, 400 MHz): δ(ppm) 5.32 (quint, 1H, O-CH(CH_2_)_2_), 4.31 (dddd, 4H, 2 × - CH_2_), 2.32 (td, 4H, 2 × -CH_2_), 1.92 (s, 6H, 2 × -CH_3_), 1.61 (quint, 4H, 2 × -CH_2_), 1.26 (m, 56H, 28 × -CH_2_), 0.88 (t, 6H, 2 × -CH_3_). (**Fig. SI 1**)

### Synthesis of 2C_18_-O(VDM) for subsequent ring-opening modification

The protocol was previously followed by our group to synthesize well-defined polymers with narrow molecular weight distributions and high end-group fidelity [53, 56]. The degree of polymerization (DP) was modulated by adjusting the monomer feed, whereby increasing the amount of VDM relative to the initiator enabled the synthesis of oligomers with target DPs of 20 and 50. A representative protocol is detailed below.

Monomer: VDM (based on DP eq.), solvent: DMF (1.0 mL), initiator: 2C_18_-Br (0.0439 g, 0.299 mmol, 1 eq.), ligand: Me_6_TREN (0.0463 mL, 0.173 mmol, 0.58 eq.), deactivating species: Cu(II)Br_2_ (0.0200 g, 0.0897 mmol, 0.30 eq.), were charged to a polymerisation flask with a magnetic stirring bar and fitted with a rubber septum. The mixture was then degassed via nitrogen sparging for 15 min, after which pre-activated copper wire was carefully added under a nitrogen blanket. The polymerisation flask was then resealed and deoxygenated for a further 5 minutes. The polymerisation was carried out at 60 °C in a temperature-controlled oil bath and stirred at 350 RPM for 5 hours. The samples were taken for proton-nuclear magnetic resonance (^1^H NMR), Gel Permeation Chromatography (GPC), and Attenuated Total Reflectance-Fourier Transform Infrared (ATR-FTIR). The samples for ^1^H NMR were diluted with CDCl_3_, while the samples for GPC were diluted with DMAc.

## 2C18-O(VDM)20

1H NMR (400.16 MHz, CDCl_3_), δ (ppm): 4.28 (s, 1H, -CH_2_-CH–O); 4.15-3.72 (m,4H, -CH_2_-C(O)H-CH_2_-) 2.52–1.26(m, 66H, 2 x CH_2_–CH_2_–C(O), -CH2–CH-); 1.08 (m, 120H, NH–C(CH_3_)_2_); 0.95 (m, 56H, 28 x -CH_2_); 0.89 (d, 6H, -O-C(O)-CH_3_-CH_3_-); 0.57 (t, 6H, 2 x -CH_3_). (**Fig. SI 2A-C**).

## 2C18-O(VDM)50

1H NMR (400.16 MHz, CDCl_3_), δ (ppm): 4.28 (s, 1H, -CH_2_-CH–O); 4.15-3.72 (m,4H, -CH_2_-C(O)H-CH_2_-) 2.52–1.26(m, 66H, 2 x CH_2_–CH_2_–C(O), -CH2–CH-); 1.08 (m, 300H, NH–C(CH_3_)_2_); 0.95 (m, 56H, 28 x -CH_2_); 0.89 (d, 6H, -O-C(O)-CH_3_-CH_3_-); 0.57 (t, 6H, 2 x -CH_3_) **(Fig. SI 2D-E**).

### Ring opening of O(VDM) with *N,N*-dimethylethylenediamine (DMEN)

Crude 2C_18_-O(VDM) oligomer solution (containing 2.99 mmol of the azlactone functionality) was added to a new vial and diluted with DMF (1 mL). Meanwhile, *N,N*-dimethylethylenediamine (0.26g, 2.99 mmol) and TEA (0.458 mL) were added to 1 mL DMF and then transferred to the oligomer solution. The solution was allowed to stir at room temperature for 2 days, and the ring-opening post-modification was confirmed by ATR-FTIR and ^1^H NMR. The DMEN-containing oligomers were purified by dialysis (MWCO 2000) against deionised water, and the solvent was removed by freeze-drying. The samples for ^1^H NMR were diluted with D_2_O.

## 2C18-O(DMEN)20

1H NMR (400.16 MHz, CDCl_3_), δ (ppm): 4.28 (s, 1H, -CH_2_-CH–O); 3.26(s, 20H, NH-CH_2_-CH_2_-); 2.56 (s, 40H, NH-CH_2_-CH_2_-); 2.52–1.26(m, 66H, 2 x CH_2_– CH_2_–C(O), -CH2–CH-); 2.25 (m, 120H, NH–C(CH_3_)_2_); 1.83 (d, 6H, -O-C(O)-CH_3_-CH_3_-); 0.95 (m, 56H, 28 x -CH_2_); 0.57 (t, 6H, 2 x -CH_3_) (**Fig. SI 3A**)

## 2C18-O(DMEN)50

1H NMR (400.16 MHz, CDCl_3_), δ (ppm): 4.28 (s, 1H, -CH_2_-CH–O); 3.26(s, 100 H, NH-CH_2_-CH_2_-); 2.56 (s, 100H, NH-CH_2_-CH_2_-); 2.52–1.26(m, 66H, 2 x CH_2_–CH_2_–C(O), -CH2–CH-); 2.25 (m, 300 H, NH–C(CH_3_)_2_); 1.83 (d, 6H, -O-C(O)-CH_3_-CH_3_-); 0.95 (m, 56H, 28 x -CH_2_); 0.57 (t, 6H, 2 x -CH_3_) (**Fig. SI 3B**)

### Ring opening of O(VDM) with *N,N*-Boc-ethylenediamine (BEDA)

Crude 2C_18_-O(VDM) oligomer solution was added to a new vial and diluted with DMF (1 mL). *N,N*-Boc-ethylenediamine (0.3g, 1.89 mmol) and TEA (0.29 mL) were added to 1 mL DMF and then transferred to the oligomer solution. The solution was allowed to stir at room temperature for 2 days, and the ring-opening post-modification was confirmed by ATR-FTIR and ^1^H NMR. The BEDA-containing oligomers were purified by dialysis (MWCO 2000) against acetone, and the solvent was removed by a vacuum oven overnight. The samples for ^1^H NMR were diluted with CDCl_3_.

## 2C18-O(BEDA)20

1H NMR (400.16 MHz, CDCl_3_), δ (ppm): 4.28 (s, 1H, -CH_2_-CH–O); 3.23 (s, 80H, NH-CH_2_-CH_2_-); 2.52–1.26(m, 66H, 2 x CH_2_–CH_2_–C(O), -CH2–CH-); 2.25 (m, 180H, C(O)-(CH_3_)_3_); 1.83 (d, 6H, -O-C(O)-CH_3_-CH_3_-); 0.95 (m, 56H, 28 x -CH_2_); 0.57 (t, 6H, 2 x -CH_3_) (**Fig. SI 3C**)

## 2C18-O(BEDA)50

1H NMR (400.16 MHz, CDCl_3_), δ (ppm): 4.98 (s, 1H, -CH_2_-CH–O); 3.23 (s, 200H, NH-CH_2_-CH_2_-); 2.52–1.26(m, 66H, 2 x CH_2_–CH_2_–C(O), -CH2–CH-); 2.25 (m, 450H, C(O)-(CH_3_)_3_); 1.83 (d, 6H, -O-C(O)-CH_3_-CH_3_-); 0.95 (m, 56H, 28 x -CH_2_); 0.57 (t, 6H, 2 x -CH_3_) (**Fig. SI 3D**).

### Deprotection of Boc-protected oligomers

A representative procedure for deprotection is detailed below.

2C_18_-O(BEDA) oligomers (100 mg) were first dissolved in DCM (1.0 mL), and TFA (1.0 mL) was then added to allow reaction overnight. The resulting deprotected oligomer solution was then evaporated to dryness under a stream of air. A further 5.0 mL of acetone was added to the mixture and evaporated to dryness. This process was repeated three times to fully remove the residual TFA and purified by dialysis (MWCO 2000) against acetone. The solvent was removed under reduced pressure by storing in a vacuum oven overnight, and the resulting product was characterized by ¹H NMR spectroscopy. The samples for ^1^H NMR were diluted with CD_3_OD.

## 2C18-O(BEDA-D)20

1H NMR (400.16 MHz, CDCl_3_, δ (ppm): 5.33 (s, 1H, C*H*–O); 4.38 (bd, 4H, O–CH(C*H*_2_OC(O))_2_); 3.54 (s, 40H, NH-C*H*_2_-CH_2_-NH_2_); 3.12 (s, 40H, NH-CH_2_-C*H*_2_-NH_2_); 1.90–1.13(m, 68H, C*H*_2_–C*H*_2_–O, C*H*_2_–C*H*); 1.42 (s, 120H, N-C(C*H*_3_)_2_); 1.25 (s, 56H, CH_3_(C*H*_2_)_14_); 1.15 (bs, 6H, OC(O)C(C*H*_3_)_2_); 0.89 (m, 6H, C*H*_3_-CH_2_) (**Fig. SI 4A**)

## 2C18-O(BEDA-D)50

1H NMR (400.16 MHz, CDCl_3_), δ (ppm): 5.33 (s, 1H, C*H*–O); 4.38 (bd, 4H, O–CH(C*H*_2_OC(O))_2_); 3.54 (s, 100H, NH-C*H*_2_-CH_2_-NH_2_); 3.12 (s, 100H, NH-CH_2_-C*H*_2_-NH_2_); 1.90–1.13(m, 158H, C*H*_2_–C*H*_2_–O, C*H*_2_–C*H*); 1.42 (s, 300H, N-C(C*H*_3_)_2_); 1.25 (s, 56H, CH_3_(C*H*_2_)_14_); 1.15 (bs, 6H, OC(O)C(C*H*_3_)_2_); 0.89 (m, 6H, C*H*_3_-CH_2_) (**Fig. SI 4B**).

### Quaternisation of synthesised DMEN-containing oligomers

2C_18_-O(DMEN) oligomer (100mg) was dissolved in DMF (10 mL), and 10 equivalents of methyl iodide (CH_3_I) per amine group were added to 1mL DMF. 2C_18_-O(DMEN) oligomer (100 mg) was dissolved in DMF (10 mL). A solution of methyl iodide (CH_3_I, 10 eq. per amine group) in DMF (1 mL) was added dropwise at room temperature, and the reaction mixture was stirred at room temperature for 4 days, resulting in complete quaternization. The product was characterized by ¹H NMR spectroscopy. Solvent and excess methyl iodide were removed by dialysis (MWCO 2000) against deionised water and then freeze-drying. The samples for ^1^H NMR were diluted with D_2_O.

## 2C18-O(DMENQ)20

1H NMR (400.16 MHz, D_2_O), δ (ppm): 5.33 (s, 1H, C*H*–O); 4.38-4.15 (bd, 4H, O–CH(C*H*_2_OC(O))_2_); 3.65 – 3.46 (d, 72H, NH-C*H*_2_-C*H*_2_); 3.16 (bs, 162H, NH-(CH_3_)_3_); 2.19–1.46 (m, 62H, C*H*_2_–C*H*, C(O)OC*H*_2_–C*H*_2_); 1.47 (bs, 108H, NH–C(CH_3_)_2_); 1.21 (s, 56H, CH_3_(C*H*_2_)_14_); 0.81 (m, 6H, C*H*_3_-CH_2_) (**Fig. SI 5A**)

## 2C18-O(DMENQ)50

1H NMR (400.16 MHz, D_2_O), 5.42 (s, 1H, C*H*–O); 4.43-4.23 (bd, 4H, O–CH(C*H*_2_OC(O))_2_); 3.76 – 3.57 (d, 176H, NH-C*H*_2_-C*H*_2_); 3.26 (bs, 396H, NH-(CH_3_)_3_); 2.41–1.47 (m, 140H, C*H*_2_–C*H*, C(O)OC*H*_2_–C*H*_2_); 1.47 (bs, 264H, NH–C(CH_3_)_2_); 1.30 (s, 56H, CH_3_(C*H*_2_)_14_); 0.89 (m, 6H, C*H*_3_-CH_2_) (**Fig. SI 5B**)

### CLO characterisation

#### 1H-nuclear magnetic resonance (^1^H NMR)

All ^1^H NMR spectra were obtained using a Bruker Avance III 400 MHz spectrometer using an external lock and referenced to the residual nondeuterated solvent. Chemical shifts (δ) were reported in parts per million (ppm). ^1^H NMR solvents (CDCl_3_, DMSO-d_6_, CD_3_OD, and D_2_O) were purchased from Cambridge Isotope Laboratories, Inc. and used as received.

### Gel permeation chromatography (GPC)

GPC analyses of oligomer samples were performed using a Shimadzu modular system comprising a DGU-20A3R degasser unit, an SIL-20A HT autosampler, a 10.0 μm bead-size guard column (50 × 7.8 mm) followed by three KF-805L columns (300 × 8 mm, bead size: 10 μm, pore size maximum: 5000 Å), an SPD-20A UV/Vis detector, and an RID-10A differential refractive-index detector. The temperature of the columns was maintained at 40 °C using a CTO-20A oven. The eluent was *N,N*-dimethylacetamide (CHROMASOLV Plus for HPLC), and the flow rate was kept at 1.0 mL/min using an LC-20AD pump. A molecular weight calibration curve was produced using commercial narrow molecular weight distribution polystyrene standards with molecular weights ranging from 500 to 2 × 10^6^ g/mol. Oligomer solutions at approx. 2 mg/mL were prepared and filtered through 0.45 μm PTFE filters before injection.

### Attenuated Total Reflectance-Fourier Transform Infrared (ATR-FTIR)

ATR-FTIR spectra were recorded using a Shimadzu IR Tracer-100 Fourier Transform Infrared spectrophotometer with an MCT detector by averaging 512 scans at a resolution of 8 cm^-1^ in the MIR region of 4000–400 cm^-1^.

### Bacterial broth microdilution assay

Bacteria were cultured in cation-adjusted Mueller Hinton broth (CaMHB) at 37 °C overnight. A sample of each culture was then diluted 40-fold in fresh broth and incubated at 37 °C for 1.5 - 3 h. The resultant mid-log phase cultures were diluted (CFU/mL measured by OD_600_), then added to each well of the compound-containing plates, giving a final cell density of 5 x 10^5^ CFU/mL and a compound concentration range of 512 to 4 µg/mL. All plates were covered and incubated at 37 °C for 18 h without shaking. MICs were determined using optical density 600 (OD_600_) after 18 h incubation by Tecan plate reader. MICs were calculated as ≥90% inhibition of bacterial growth compared to the positive control wells. Analysis was performed using Pipeline Pilot/Python script.

### Fungal broth microdilution assay

Fungal strains were cultured for 2 days on Potato Dextrose Agar (PDA) agar at 35 °C. A yeast suspension of 1 × 10^6^ to 5 × 10^6^ CFU/mL (as determined by OD_530_) was prepared from several colonies. The suspension was subsequently diluted and added to each well of the compound-containing plates, giving a final cell density of 2.5 × 10^3^ CFU/mL and a compound concentration range of 512 to 4 µg/mL. All plates were covered and incubated at 35 °C for 36 h without shaking. MICs were determined by optical density after 36 h incubation using the Biotek Synergy HTx Plate reader. For *Candida albicans,* OD_630_ was recorded, and for *Cryptococcus neoformans,* OD_600-570_ was recorded after the addition of 0.01% resazurin. MICs were calculated as ≥90% inhibition of fungal growth compared to the positive control wells. Analysis was performed using Pipeline Pilot/Python script.

### Cell viability assay

HEK293 cells were suspended in DMEM medium (Gibco; 11330332), supplemented with 10% FBS (GE; SH30084.03) and 100 U/mL each of penicillin and streptomycin (Invitrogen; 15070063), and were added into the test plates at 5000 cells per well in a volume of 48 µL. The final concentration range was 4 – 512 µg/mL. The cell plates were incubated for 20 h at 37 °C, 5% CO_2_. After incubation, 5 µL of 100 µM resazurin (Sigma; R7017) in PBS was added to each well for a final concentration of 11 µM. The plates were then incubated for 3 h at 37 °C, 5% CO_2_. The fluorescence intensity was read using the TECAN Infinite M1000 PRO with excitation/emission at 560/590 nm, respectively.

Cell viability was calculated using the following equation:

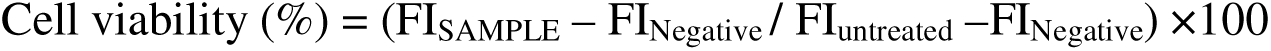

Using nonlinear regression analysis of log(concentration) *vs* normalised cell viability, using variable fitting, CC_50_ (concentration at 50% cell viability) was calculated. In addition, the maximum percentage of cell inhibition was reported. Any value with > indicates a sample with no inhibition or CC_50_ above the maximum tested concentration.

### Haemolysis assay

Human whole blood (10 mL/tube) was washed 2-3 times in 3 volumes of 0.9% NaCl (Baxter; AHF7124), with centrifugation for 10 min between washes, at 500 g, with reduced deceleration. A cell count was performed using a Neubauer haemocytometer, and then cells were diluted to 1 × 10^8^/mL in 0.9% NaCl. Diluted cells (48 µL/well) were added to the compound-containing PP plates. Plates were sealed and then placed on a plate shaker for 10 min before being incubated for 1 h at 37 °C without shaking.

After incubation plates were centrifuged at 1000 g for 10 min, to pellet cells and debris, and then 25 µL of the supernatant was transferred into a 384-well flat-bottom polystyrene (PS) plate (Corning; Cat. No. 3680). Following transfer, absorbance (Abs) was read at 405 nm using a Tecan M1000 Pro monochromator plate reader. Analysis was performed using Pipeline Pilot/Python script. The percentage of haemolysis was calculated using the following equation:

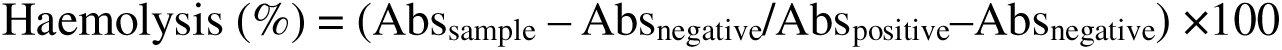

Using nonlinear regression analysis of log(concentration) vs normalised haemolysis, using variable fitting, HC_10_ and HC_50_ (concentration at 10 and 50% haemolysis, respectively) were calculated. In addition, the maximum percentage of haemolysis is reported. Any value with ‘>’ indicates samples with no haemolytic activity or a HC_10_ or HC_50_ above the maximum tested concentration.

## Results and Discussion

A library of 1,2-*O*-dioctadecyl (2C_18_)-terminated cationic lipidated oligomers with the targeted degree of polymerisation (DP) of 20 and 50 was synthesised using a VDM-functional oligomeric platform employing a two-step, single-pot process (**Scheme 1**).

### Synthesis of precursor lipidated oligomers

First, 2C_18_-containing lipidated oligomers were synthesised using an optimised Cu(0)-mediated RDRP conditions (2C_18_-Br : CuBr_2_ : Me_6_Tren = 1 : 0.3 : 0.58). The targeted DP was achieved via modifying the ratio of VDM monomer to initiator (20 eq. and 50 eq., respectively). DMF was used as an effective polar solvent to provide an efficient disproportionation for controlled polymerisation of both hydrophobic and hydrophilic materials [62, 63]. The successful polymerisation of oligomers, with well-controlled molecular weight and low dispersity (*Ð*), was confirmed by ^1^H NMR and GPC. In all cases, relatively low *Ð* (1.2 to 1.3) and acceptable quasi-full conversion (∼90%) were achieved in five hours (**Table SI 1 and Fig. SI 6**). Furthermore, characterisation by ATR-FTIR showed that the peak at 1813 cm^-1^ (corresponding to the carbonyl groups [C=O] from oxazolone ring) was preserved throughout the polymerisation (**Fig. 1**).

**Figure 1.**
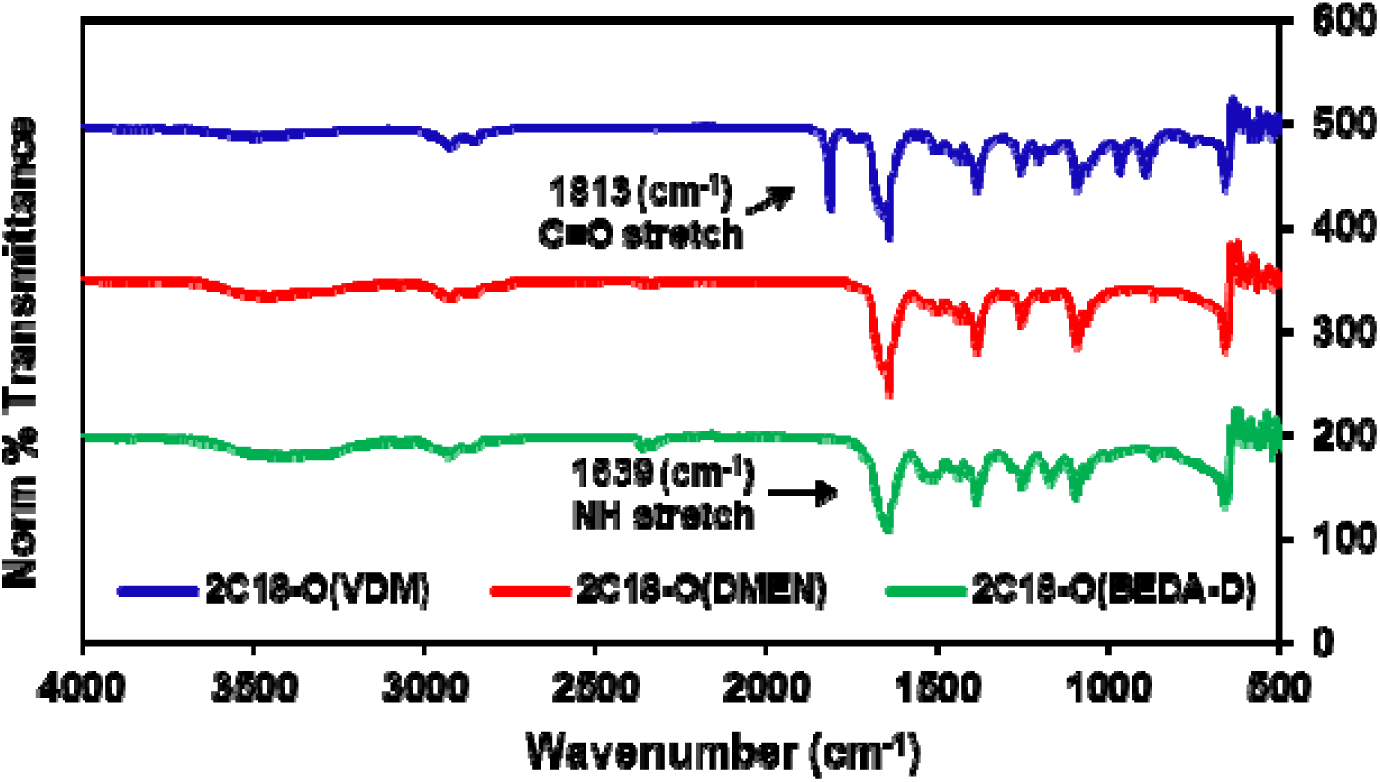
ATR-FTIR spectra of 2C_18_-terminated oligomers: unreacted O(VDM) (royal blue), and the final ring-opened oligomers containing: DMEN (red); BEDA-D (green), 512 scans at a resolution of 8 cm^-1^.

**Scheme 1.**
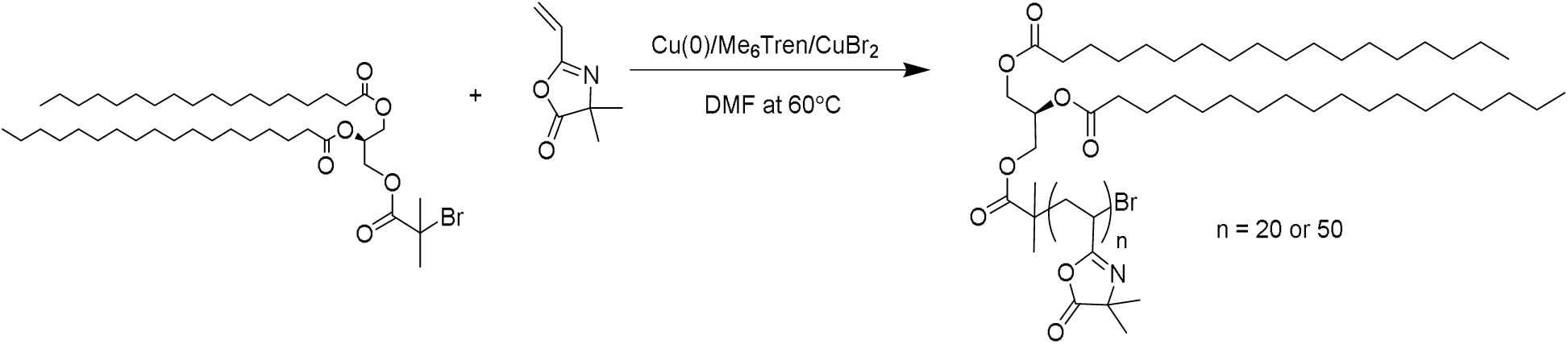
Cu(0)-mediated polymerisation of 2-vinyl-4,4-dimethyl-5-oxazolone (VDM) using (R)-3-((2-bromo-2-methylpropanoyl)oxy)propane-1,2-diyl dioctadecanoate initiator (2C_18_-Br); *n* = 20, 50.

### Ring opening of oligo(VDM)

All 2C_18_ functionalised VDM oligomers were subsequently ring-opened with two commercially available reactive amines, including a Boc-protected primary amine (BEDA) and a tertiary amine (DMEN) (**Scheme 2**) to give a final library of 4 cationic lipidated oligomers (CLOs). These amine compounds were expected to exhibit native antibacterial and antifungal activities and after deprotection of the Boc-protected groups. These specific precursors of cationic groups were chosen, as they have been previously reported to exhibit antimicrobial activity [48, 60–63]. CLOs with primary amine pendant groups are essentially lysine mimics and have been reported to have antimicrobial activities previously [64]. Specifically, *N, N*-dimethylethylenediamine (DMEN) and *N*-Boc-ethylenediamine (BEDA) were found to be the most suitable cationic groups that exhibited superior antimicrobial activities in a range of functional amines. ^1^H NMR and FTIR spectra were also taken after ring opening to confirm successful ring opening, and the molecular weight and polydispersity are presented in **Table 1**.

**Table 1.** Characterisation of ring-opened oligomers by ^1^H NMR spectroscopy and GPC.

| Ring opened with | CLOs | $^1\text{H}$ NMR | | | GPC | |
| --- | --- | --- | --- | --- | --- | --- |
| | | Conv.<sup>a</sup> (%) | DP<sup>a</sup> | $M_{n,\text{th}}^b$<br>(g mol<sup>-1</sup>) | $M_n^c$<br>(g mol<sup>-1</sup>) | $\bar{D}^c$ |
| *N, N*-Dimethylethylenediamine | 2C<sub>18</sub>-O(DMEN)<sub>20</sub> | ~96 | 20 | 5400 | -<sup>d</sup> | -<sup>d</sup> |
| *N, N*-Dimethylethylenediamine | 2C<sub>18</sub>-O(DMENQ)<sub>20</sub> | ~94 | 18 | 5700 | -<sup>d</sup> | -<sup>d</sup> |
| *N*-Boc-ethylenediamine | 2C<sub>18</sub>-O(BEDA)<sub>20</sub> | ~92 | 18 | 6841 | 7131 | 1.21 |
| *N, N*-Dimethylethylenediamine | 2C<sub>18</sub>-O(DMENQ)<sub>50</sub> | ~82 | 44 | 12219 | -<sup>d</sup> | -<sup>d</sup> |
| *N, N*-Dimethylethylenediamine | 2C<sub>18</sub>-O(DMEN)<sub>50</sub> | ~83 | 42 | 12969 | -<sup>d</sup> | -<sup>d</sup> |
| *N*-Boc-ethylenediamine | 2C<sub>18</sub>-O(BEDA)<sub>50</sub> | ~93 | 43 | 15822 | 14621 | 1.13 |

<sup>a</sup> Conversion (%), DP (degree of polymerisation) values were determined by $^1\text{H}$ NMR peak integration analysis. <sup>b</sup> $M_{n,\text{th}}$ was calculated for the homo-oligomer based on the conversion, and the $M_{n,\text{th}}$ for the ring-opened oligomer was calculated by assuming full ring-opening of the oligomer by the amine added as reported in the ESI. <sup>c</sup> $M_n$ and $\bar{D}$ were determined by GPC analysis in DMAc against polystyrene standards. <sup>d</sup> GPC analysis could not be obtained for oligomers containing DMEN.

**Scheme 2.**
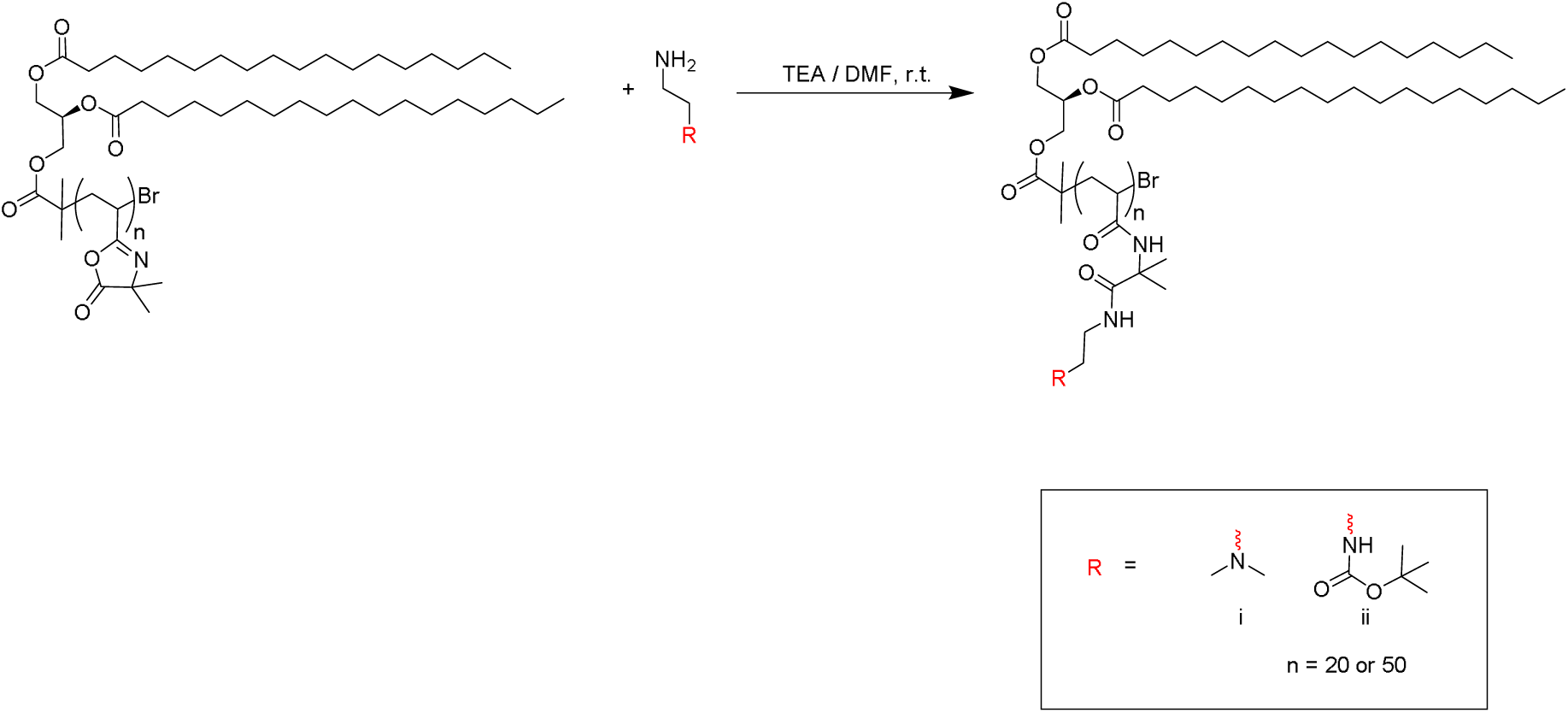
Representative scheme of ring-opening of 2C_18_-O(VDM) with two different functionalised reactive amine moieties, i.e., (i) *N, N*-dimethylethylenediamine and (ii) *N*-Boc-ethylenediamine.

The successful ring opening of the oxazolone moiety was confirmed by ATR-FTIR spectroscopy, as evidenced by the disappearance of the characteristic absorption band at ∼1813 cm^-1^, attributable to the C=O stretching vibration of the oxazolone ring, and the concomitant emergence of a strong band at ∼1639 cm^-1^, corresponding to the amide carbonyl vibrations of the two amide functionalities formed upon ring opening (**Fig. 1**).

It was observed that the oligomers ring-opened with DMEN could not be detected via GPC. Literature suggested that the addition of small amounts of triethylamine (TEA) may enhance peak resolution; however, under our experimental conditions, no such improvement was observed [53, 65]. This behaviour is likely attributable to the strong cationic nature of the ring-opened oligomers, which can promote electrostatic interactions with the GPC column and result in poor elution, consistent with observations for related cationic systems. Therefore, ^1^H NMR was relied on to indicate the modification via the appearance of new peaks corresponding to the incorporated pendant groups. The ^1^H NMR spectra of DMEN-functionalized oligomers exhibited additional resonances attributable to the incorporated DMEN groups at ∼3.37 ppm corresponding to the -NH-C**H_2_**-C**H_2_**-N- protons (**Fig. SI 3A & B**). Similarly, the ^1^H NMR spectra of BEDA-containing oligomers displayed a combined resonance at ∼3.40 ppm, corresponding to –NH–C**H_2_**–C**H_2_**–NH– protons of the incorporated BEDA groups. (**Fig. SI 3C & D**). The completeness of ring opening was assessed by ¹H NMR by comparing the disappearance of the oxazolone signal with the appearance of new resonances corresponding to the modified oligomer, using a nonreactive lipid signal as an internal reference. Quantitative conversion was confirmed when the integration ratio matched the expected degree of polymerization.

### Deprotection of Boc-protecting groups

The ring-opened BEDA containing oligomers were then deprotected to reveal primary amine functional groups. The deprotection of the Boc-protected BEDA was achieved via reaction with an excess of trifluoroacetic acid (TFA), and the primary amine group was thus revealed (**Scheme 3**).

**Scheme 3.**
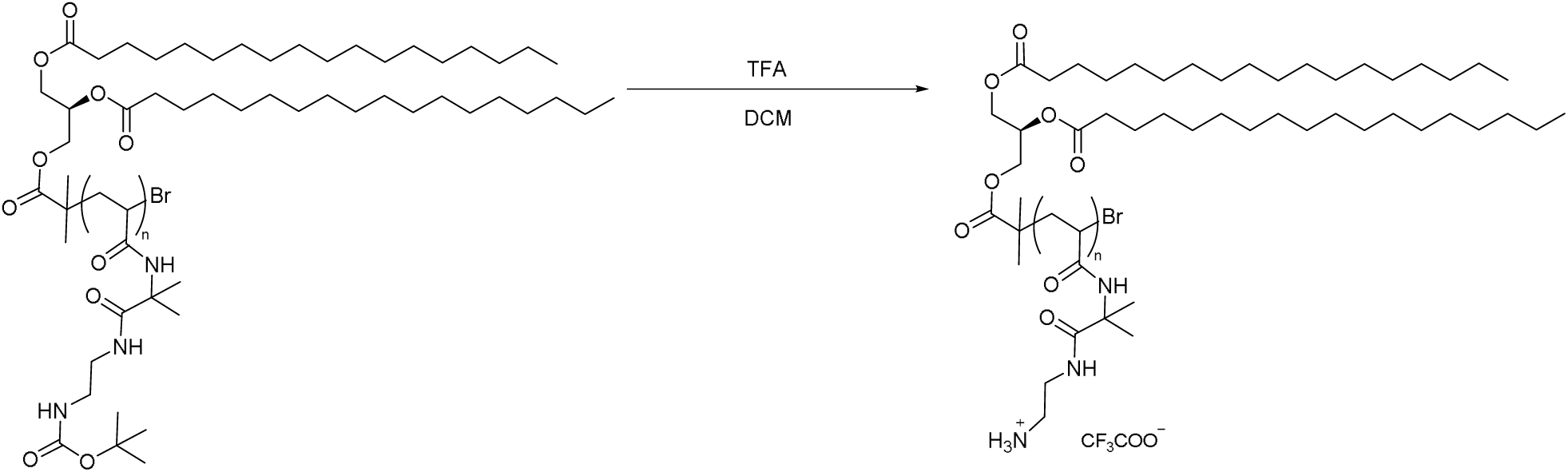
Representative scheme depicting deprotection of the Boc-groups utilising trifluoroacetic acid (TFA), where *n* = 20, 50

^1^ H NMR was used to confirm the full deprotection (**Fig. SI 4**). Specifically, in contrast to the pre-modification spectrum, the peak around ∼2.2 ppm (corresponding to tert-butyl hydrogen (-C-C**H_3_**-C**H_3_**-C**H_3_**-)) disappeared. Moreover, ^1^H NMR spectra revealed that the peak at ∼3.40 ppm was split into two individual peaks, which represented the -C**H_2_**-C**H_2_**- structure in the side chain. Because the post-modification reaction did not alter the oligomer backbone, the DP was expected to remain unchanged after deprotection. Accordingly, in the ^1^H NMR spectra, the integration ratio of unchanged backbone or lipid resonances relative to the original oligomer signals remained constant. In contrast, signals associated with the protecting groups were lost. However, the deprotected oligomers could not be characterized by GPC because their strong cationic character resulted in poor elution. The experimental molecular weight was thus calculated based on the corresponding peaks in the ^1^H NMR, noting that the equivalent ratio of TFA salt (CF_3_COO^-^) was included in the calculation (**Table 1**).

### Quaternisation of tertiary amine groups

To broaden the antimicrobial potential of CLOs, our group previously introduced quaternary ammonium cations (QACs) into CLOs, which, in this previous work, was found to enhance antimicrobial activity [66]. DMEN-containing CLOs were exposed to an excess of iodomethane to convert the tertiary amine group to a quaternary ammonium group (**Scheme 4**). ^1^H NMR revealed essentially the full conversion of the tertiary amine groups, via a downfield shift of peaks corresponding to -C**H_2_**-C**H_2_**- protons from ∼2.5 - 3.4ppm to ∼3.4 - 3.5 ppm respectively. The success of quaternisation of tertiary amino oligomers was confirmed via the presence of methyl hydrogens surrounding the nitrogen atom at ∼3.2 ppm, and the DP determined from the peak integrals (at I_3.2 ppm_) was identical to the pre-methylated oligomers (**Fig. SI 5**). Due to their strong cationic character, the quaternised DMEN-oligomers could not be accurately characterised by GPC, as discussed previously. Consequently, the experimental molecular weight was estimated by incorporating the molecular mass of the corresponding iodide counterion.

**Scheme 4.**
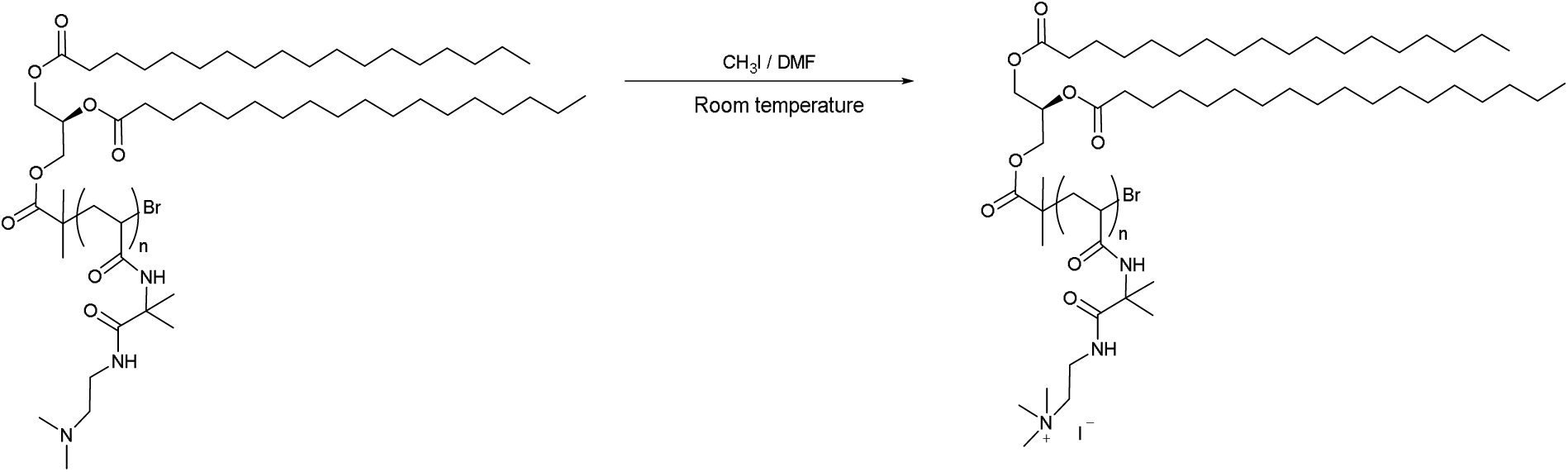
Representative scheme of quaternisation of tertiary amine groups using methyl iodide, where *n* = 20, 50

### Minimum Inhibitory Concentration (MIC) for various pathogens

The library of 6 CLOs was investigated against a panel of WHO-priority bacterial and fungal pathogens [Fungi; *Candida albicans* ATCC 90028, *Cryptococcus neoformans* ATCC 208821, Gram-negative; *Escherichia coli* ATCC 25922, *Klebsiella pneumoniae* ATCC 700603, *Acinetobacter baumannii* ATCC 19606, *Pseudomonas aeruginosa* ATCC 27853, Gram-positive; methicillin-resistant *Staphylococcus aureus* (MRSA) ATCC 43300]. The CLOs exhibited superior antibacterial activity than antifungal activity (see **Table 2** *vs.* **Table SI 2**).

**Table 2.** Antibacterial testing of CLOs against Gram-positive strain; *Methicillin-resistant S. aureus* (MRSA) ATCC 43300, and Gram-negative strains; *E. coli* ATCC 25922, *K. pneumoniae* ATCC 700603, *A. baumannii* ATCC 19606, and *P. aeruginosa* ATCC 27853.

| CLOs | MIC ($\mu\text{g mL}^{-1}$) | | | | |
| --- | --- | --- | --- | --- | --- |
|  | *E. coli* | *K. pneumoniae* | *A. baumannii* | *P. aeruginosa* | MRSA |
| 2C<sub>18</sub>-O(DMEN)<sub>20</sub> | >512 | >512 | >512 | >512 | $\leq 4$ |
| 2C<sub>18</sub>-O(DMENQ)<sub>20</sub> | >512 | >512 | >512 | >512 | $\leq 4$ |
| 2C<sub>18</sub>-O(BEDA)<sub>20</sub> | >512 | >512 | 64 | >512 | $\leq 4$ |
| 2C<sub>18</sub>-O(DMENQ)<sub>50</sub> | >512 | >512 | >512 | >512 | $\leq 4$ |
| 2C<sub>18</sub>-O(DMEN)<sub>50</sub> | >512 | >512 | $\leq 4$ | >512 | >512 |
| 2C<sub>18</sub>-O(BEDA)<sub>50</sub> | $\leq 4$ | >512 | $\leq 4$ | 32* | $\leq 4$ |

\*MIC values for this compound showed variability across replicate experiments

All CLOs showed no detectable activity against *K. pneumoniae* and *E. coli* (MIC > 512 µg mL^-1^) **(Table 2)**. In contrast, several CLOs displayed strong antibacterial activity against *S. aureus* (MRSA) and *A. baumannii* (MIC ≤ 4 µg mL^-1^). With the exception of 2C_18_-O(DMEN)_50_, all CLOs exhibited potent activity against Gram-positive MRSA. The synthesized CLOs shared a common terminal group (2C_18_) but differed in pendant amine functionalities: DMEN (tertiary amine), BEDA (primary amine), and DMENQ (quaternary ammonium). By maintaining the end-group constant while varying the degree of polymerization (DP), it was evident that the BEDA series displayed the broadest antibacterial spectrum. Notably, increasing the DP from 20 to 50 significantly improved the activity of BEDA-based CLOs against *A. baumannii*, reducing the MIC from 64 µg mL^-1^ to ≤ 4 µg mL^-1^ **(Table 2)**. Interestingly, compared to oligomers with primary amine pendants and ethyl (C_2_), dodecyl (C_12_) or diglyceride (2C_12_) end-groups, as previously reported by our group, using a more hydrophobic end group was likely to change the antimicrobial profile from favouring antifungal activity to antibacterial activity [53].

The antifungal activity of the CLOs was also evaluated against fungal pathogens, *Candida albicans* ATCC 90028 and *Cryptococcus neoformans* H99 (ATCC 208821). All CLOs exhibited minimal activity against *C. albicans* (MIC > 512 µg mL^-1^), indicating limited efficacy toward this species. In contrast, several members of the DMEN series, including 2C_18_-O(DMEN)_20_, 2C_18_-O(DMEN)_50_, and 2C_18_-O(DMENQ)_50_, demonstrated potent activity against *C. neoformans* (MIC ≤ 4 µg mL^-1^), comparable to that of established antifungal agents. The quaternary analogue 2C_18_-O(DMENQ)_20_ showed moderate inhibition (MIC =128 µg mL^-1^). These results suggest that increased oligomer length and quaternisation enhance antifungal potency, likely by promoting stronger interactions with and disruption of the fungal cell membrane [67]. Considering that fungal cell membranes contain ergosterol, a key component responsible for maintaining membrane integrity, whereas bacterial membranes lack sterols and instead comprise structural elements such as peptidoglycan [68], this fundamental difference in membrane composition likely underlies the greater antibacterial activity observed for these polymers [26, 69, 70]. However, further research and comparative studies are required to elucidate the specific molecular basis underlying the differential efficacy of 2C_18_-terminated CLOs as antibacterial rather than antifungal agents.

### Haemolysis assay

The hemocompatibility of the CLOs was assessed through haemolysis assays against human red blood cells. HC_50_ is defined as the oligomer concentration that causes 50% haemolysis, which is commonly used to reflect the haemotoxicity of an agent [71, 72]. All CLOs exhibited HC_50_ values greater than 512 µg/mL^-1^, demonstrating negligible haemolytic activity even at the highest concentrations tested [42]. (**Table 3**)

**Table 3.**
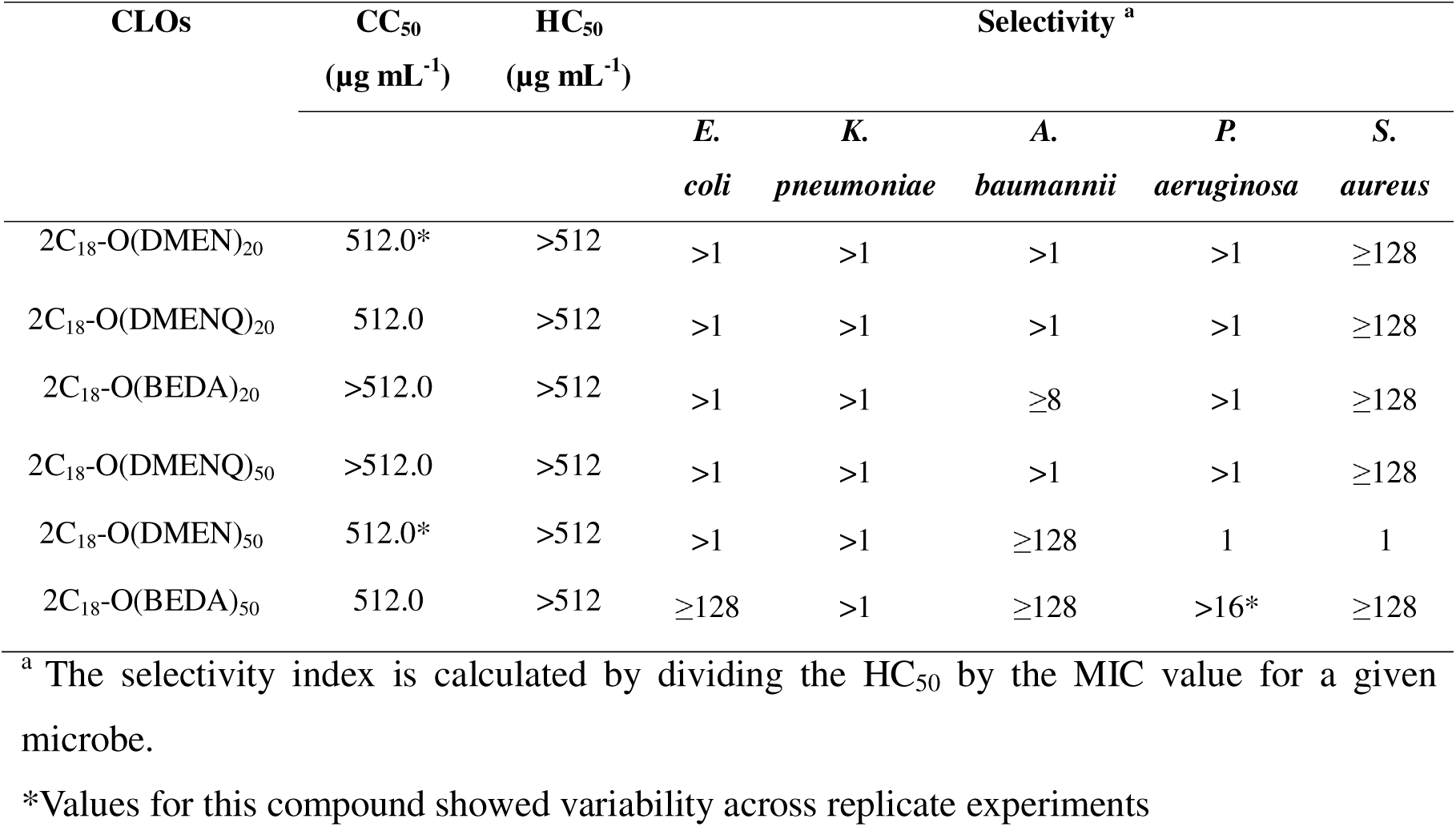
Selectivity index values for the oligomer library against Gram-positive and Gram-negative bacterial strains.

| CLOs | CC<sub>50</sub><br>($\mu\text{g mL}^{-1}$) | HC<sub>50</sub><br>($\mu\text{g mL}^{-1}$) | Selectivity <sup>a</sup> | | | | |
| --- | --- | --- | --- | --- | --- | --- | --- |
|  |  |  | *E. coli* | *K. pneumoniae* | *A. baumannii* | *P. aeruginosa* | *S. aureus* |
| 2C<sub>18</sub>-O(DMEN)<sub>20</sub> | 512.0* | >512 | >1 | >1 | >1 | >1 | ≥128 |
| 2C<sub>18</sub>-O(DMENQ)<sub>20</sub> | 512.0 | >512 | >1 | >1 | >1 | >1 | ≥128 |
| 2C<sub>18</sub>-O(BEDA)<sub>20</sub> | >512.0 | >512 | >1 | >1 | ≥8 | >1 | ≥128 |
| 2C<sub>18</sub>-O(DMENQ)<sub>50</sub> | >512.0 | >512 | >1 | >1 | >1 | >1 | ≥128 |
| 2C<sub>18</sub>-O(DMEN)<sub>50</sub> | 512.0* | >512 | >1 | >1 | ≥128 | 1 | 1 |
| 2C<sub>18</sub>-O(BEDA)<sub>50</sub> | 512.0 | >512 | ≥128 | >1 | ≥128 | >16* | ≥128 |

<sup>a</sup> The selectivity index is calculated by dividing the HC<sub>50</sub> by the MIC value for a given microbe.
\*Values for this compound showed variability across replicate experiments

### Cell viability

Mammalian cytotoxicity (CC_50_) of the CLOs was evaluated using a cell viability assay in HEK293 cells [73]. All CLOs exhibited low cytotoxicity, with CC_50_ values generally ≥ 512 µg mL^-1^. Minor variability was observed for 2C_18_-O(DMEN)_20_ and 2C_18_-O(DMEN)_50_ (**Table 3**). The consistently high CC_50_ values indicate that these CLOs are well tolerated by mammalian cells under the tested conditions, thereby providing a broad therapeutic window in which antimicrobial activity can be exerted without significant host cell toxicity.

### Selectivity

The therapeutic potential of CLOs was further assessed by examining their selectivity, which was calculated as the ratio of hemolytic activity (expressed as the HC_50_) to the antimicrobial minimum inhibitory concentrations (MICs). Against Gram-negative bacteria (*E. coli*, *K. pneumoniae*, *A. baumannii*, and *P. aeruginosa*), the oligomers generally exhibited limited antimicrobial activity, reflected by low or modest selectivity indices. Notably, however, 2C_18_-O(BEDA)_50_ showed enhanced activity and selectivity against *E. coli* and *A. baumannii* (SI ≥ 128). In contrast, consistently high selectivity was observed against the Gram-positive pathogen S. aureus across the library (SI ≥ 128). These results support the finding proposed by Togahi *et al.* that longer chain fatty acid exhibits greater antibacterial activity against *S. aureus*, with octadecanol specifically showing the highest selectivity (MIC > 512 µg mL^-1^) against the pathogen [74]. Such differential activity emphasizes the importance of optimizing oligomer composition and degree of polymerization to achieve a balance between potency and safety and positions these CLOs as promising candidates for further therapeutic development (see **Table 3** *vs* **Table SI 3**).

## Conclusion

We report the controlled synthesis of antimicrobial peptide (AMP)-mimicking cationic lipidated oligomers (CLOs) via Cu(0)-mediated RDRP using 2C_18_-Br as the initiator. A library of VDM-derived oligomers with target degrees of polymerization of 20 or 50 was successfully synthesised, and subsequently functionalized via nucleophilic ring opening to yield primary (BEDA) and tertiary (DMEN) amines. The library was further expanded via quaternarisation of the tertiary amine pendant groups (DMEN), yielding DMENQ. Antimicrobial testing revealed that all CLOs except one were inactive against Gram-negative *E. coli* and *K. pneumoniae* (MIC > 512 µg mL^-1^), and two showed high activity against *A. baumannii* (MIC ≤4 µg mL^-1^). Moreover, several CLOs showed potent activity against Gram-positive *S. aureus* (MRSA) (MIC ≤ 4 µg mL^-1^). The BEDA series demonstrated the broadest antibacterial spectrum, with increased DP markedly enhancing activity toward *A. baumannii* (from 64 µg mL^-1^ at DP 20 to ≤ 4 µg mL^-1^ at DP 50). In contrast, antifungal assays indicated limited activity against *C. albicans* but notable potency of the DMEN and DMENQ series against *C. neoformans* (MIC ≤ 4 µg mL^-1^). All CLOs displayed excellent hemocompatibility (HC_50_ > 512 µg mL^-1^) and low cytotoxicity toward HEK293 cells (CC_50_ ≥ 512 µg mL^-1^). Collectively, these findings highlight the influence of end-group lipidation, cationic functionality, and polymer length on antibacterial selectivity and establish 2C_18_-terminated CLOs as promising scaffolds for developing biocompatible antimicrobial materials.

## Data availability

The data supporting this article have been included as part of the ESI. Further data will be made available upon reasonable request from reviewers.

## CRediT authorship contribution statement

**Qiaomingxuan Liu:** Methodology, Software, Formal analysis, Investigation, Writing – original draft & editing.

**Shiwei Zhang:** Methodology, Software, Formal analysis, Investigation. **Margaret Pywell:** Methodology, Software, Formal analysis, Investigation. **Alysha G. Elliott:** Resources, Data curation, Validation, Formal analysis. **Holly Floyd:** Resources, Data curation, Validation, Formal analysis.

**Johannes Zuegg:** Resources, Data curation, Validation, Formal analysis.

**Jessica R. Tait:** Supervision, Validation, Writing – review & editing

**John F. Quinn:** Funding acquisition, Supervision, Validation, Writing – review & editing **Michael R. Whittaker:** Conceptualization, Methodology, Funding acquisition, Supervision, Validation, Writing – review & editing

**Muhammad Bilal Hassan Mahboob:** Conceptualization, Methodology, Supervision, Software, Formal analysis, Investigation, Writing – review & editing.

**Cornelia B. Landersdorfer:** Conceptualization, Methodology, Funding acquisition, Supervision, Validation, Writing – review & editing

## Acknowledgement

This work was supported by the Australian Research Council (DP200102829) and the Monash Institute of Pharmacy and Pharmaceutical Sciences (MIPS). J. F. Q. is grateful for an ARC Future Fellowship (FT170100144).

