## Supplementary material for "Harnessing Diacylglycerol-Terminated Cationic Oligomers for Next-Generation Antibacterial Therapeutics": SI File

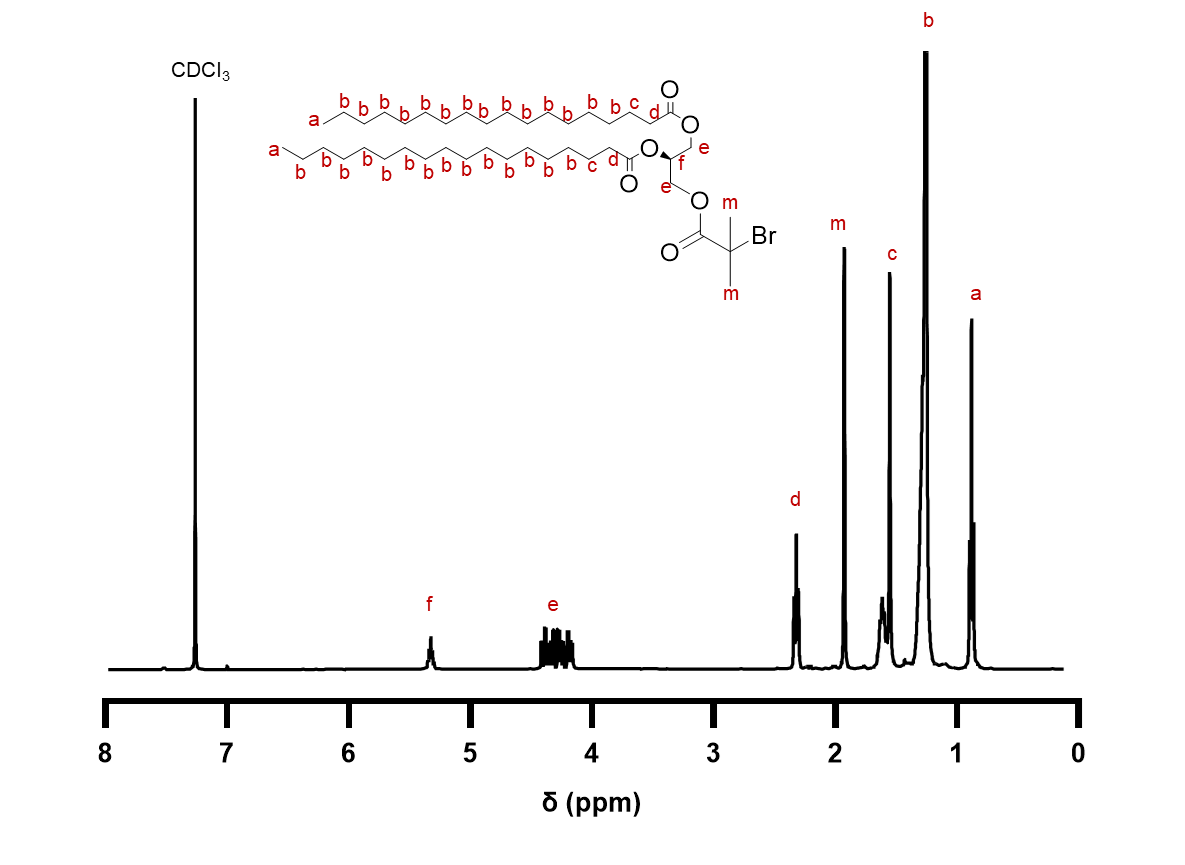

**Figure SI 1.** ^1^H NMR (400MHz, CDCl_3_) spectrum of 2C_18_-Br initiator.

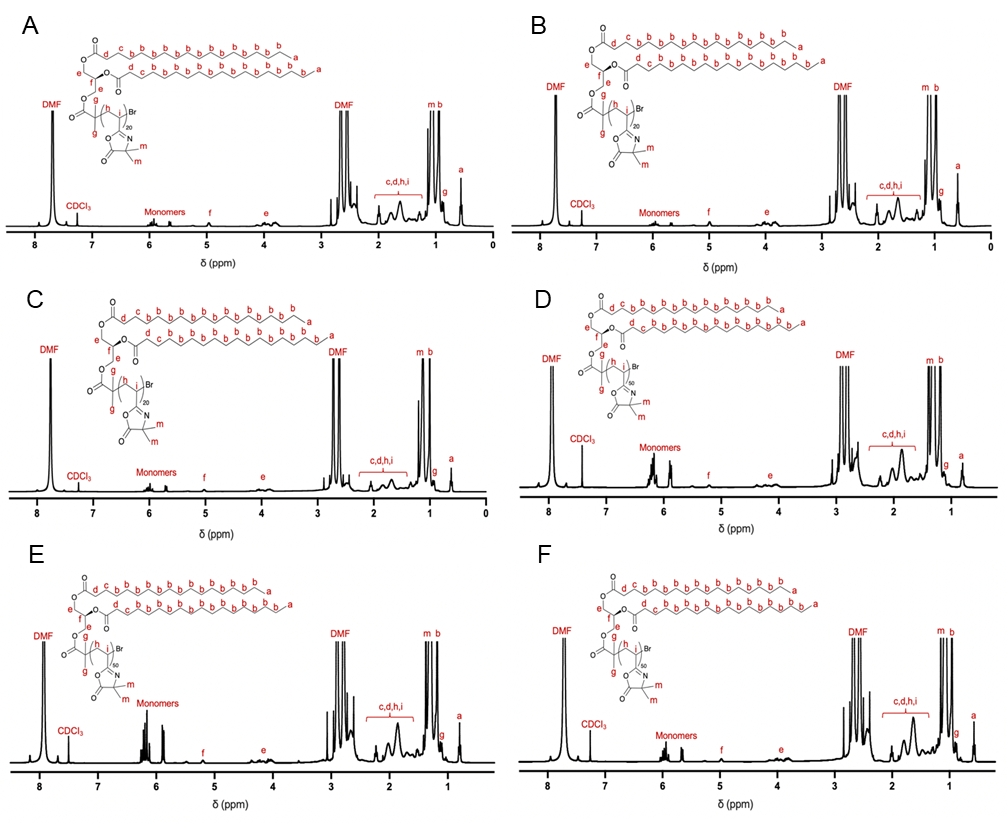

**Figure SI 2**. ^1^H NMR (400MHz, CDCl_3_) spectra of precursor O(VDM) oligomers: **A)** 2C_18_-O(VDM)_20_-1. **B)** 2C_18_-O(VDM)_20_-2. **C)** 2C_18_-O(VDM)_20_-3. **D)** 2C_18_-O(VDM)_50_-1. **E)** 2C_18_-O(VDM)_50_-2. and **F)** 2C_18_-O(VDM)_50_-3.

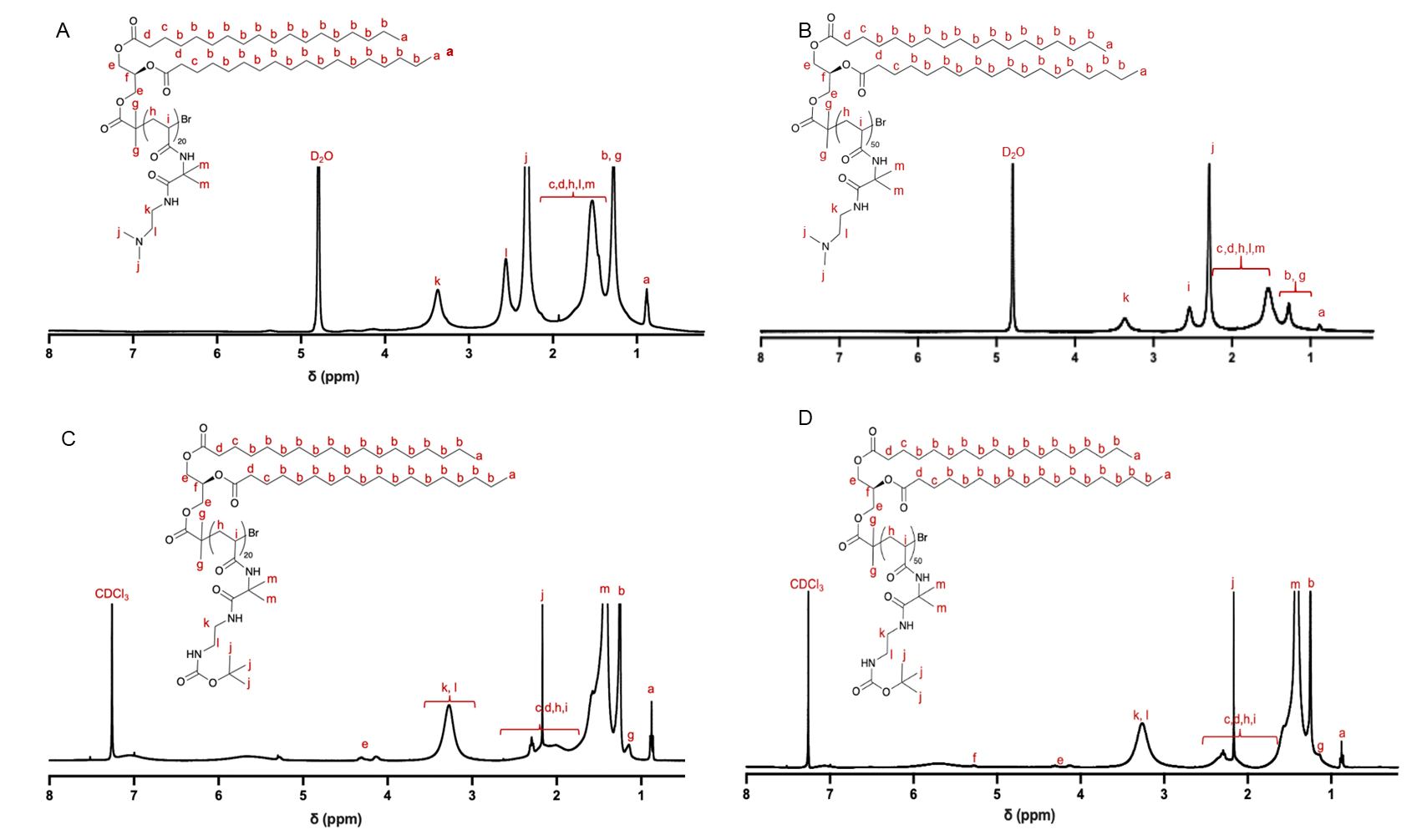

**Figure SI 3**. ^1^H NMR [400MHz, CDCl_3_ (A,B), D_2_O (C,D)]) spectra of ring-opened O(VDM) oligomers: **A)** 2C_18_-O(DMEN)_20_. **B)** 2C_18_-O(DMEN)_50_. **C)** 2C_18_-O(BEDA)_20_. **D)** 2C_18_-O(BEDA)_50_.

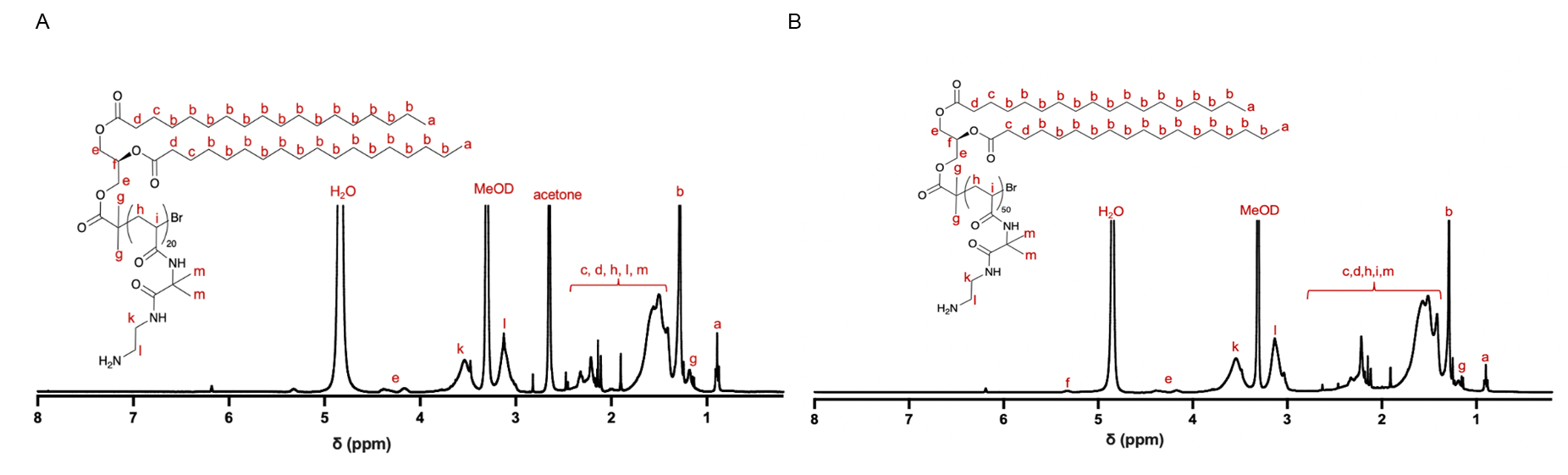

**Figure SI 4**. ^1^H NMR (400MHz, MeOD) spectra of deprotected O(BEDA) oligomers: A) 2C_18_-O(BEDA-D)_20_. B) 2C_18_-O(BEDA-D)_50_.

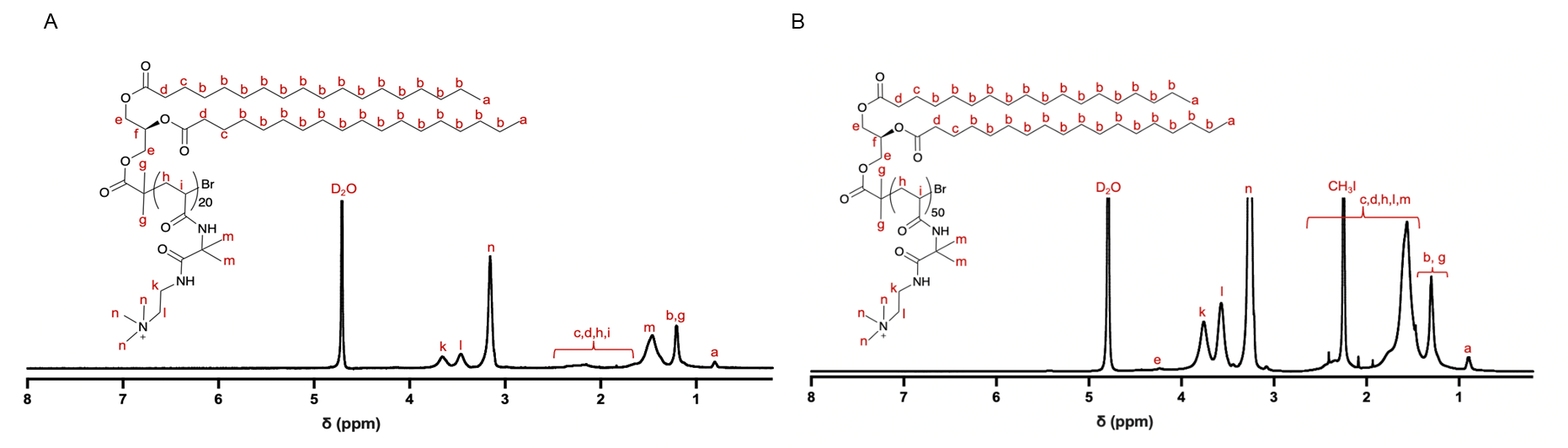

**Figure SI 5**. ^1^H NMR (400MHz, D_2_O) spectra of quaternised O(DMEN) oligomers: A) 2C_18_-O(DMENQ)_20_. B) 2C_18_-O(DMENQ)_50_.

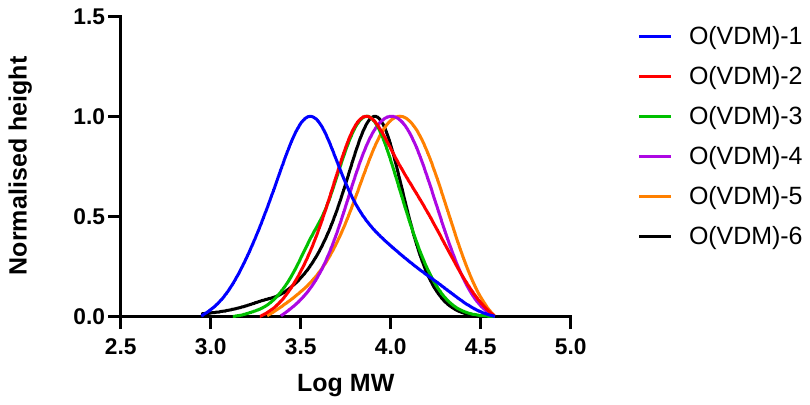

**Figure SI 6**. Molecular weight distribution of 6 repeated trials of 2C_18_-O(VDM); GPC was conducted in DMAc.

**Table SI 1.** Characterization of oligomers from Cu-mediated polymerisation by ^1^H NMR spectroscopy and GPC

| **Sr No** | **Oligomers** | **^1^H NMR** | | | | |  | **GPC** | | |
| --- | --- | --- | --- | --- | --- | --- | --- | --- | --- | --- |
|  |  | **Conversion %^a^** | **DP** | **M_n,th_^b^** |  | **M_n_^c^** | | | **Ð^c^** | |
| 1 | 2C_18_-O(VDM)_20_ | 96 | 20 | 3557 |  | 3300 | | | | 1.24 |
| 2 | 2C_18_-O(VDM)_20_ | 94 | 18 | 3279 |  | 3800 | | | | 1.35 |
| 3 | 2C_18_-O(VDM)_20_ | 92 | 18 | 3250 |  | 3600 | | | | 1.28 |
| 4 | 2C_18_-O(VDM)_50_ | 82 | 44 | 6869 |  | 6400 | | | | 1.32 |
| 5 | 2C_18_-O(VDM)_50_ | 83 | 42 | 6618 |  | 5900 | | | | 1.26 |
| 6 | 2C_18_-O(VDM)_50_ | 93 | 43 | 6757 |  | 6500 | | | | 1.33 |

a) *Conversion (%) after 5 h reaction with 2C_18_-Br: CuBr_2_: Me_6_TREN (1 0.30: 0.58), DP (degree of polymerisation) values were determined by ^1^H NMR peak integration analysis. b) M_n,th_ was calculated for the homo-oligomer based on the conversion. c) Mn and *Ð* were determined by GPC analysis in DMAc against polystyrene standards.

**Table S2. Antifungal testing of oligomer compounds against *C. albicans* ATCC 90028 and *C. neoformans* H99 ATCC 208821**

| **CLOs** | **MIC (µg mL^-1^)** |
| --- | --- |

|  | ***C. albicans*** | ***C. neoformans*** |
| --- | --- | --- |
| 2C_18_-O(DMEN)_20_ | >512 | ≤ 4 |
| 2C_18_-O(DMENQ)_20_ | >512 | 128* |
| 2C_18_-O(BEDA)_20_ | >512 | 512* |
| 2C_18_-O(DMENQ)_50_ | >512 | ≤ 4 |
| 2C_18_-O(DMEN)_50_ | 512 | ≤ 4 |
| 2C_18_-O(BEDA)_50_ | >512 | 512 |

*Values for this compound showed variability across replicate experiments

**Table S3.** **Selectivity index values for the oligomer library against *C. albicans* ATCC 90028 and *C. neoformans* H99 ATCC 208821**

| **CLOs** | **CC_50_ (µg mL^-1^)** | **HC_50_ (µg mL^-1^)** | **Selectivity** | |
| --- | --- | --- | --- | --- |
|  |  |  | ***C. albicans*** | ***C. neoformans*** |
| 2C_18_-O(DMEN)_20_ | 512.0* | >512 | >1 | ≥128 |
| 2C_18_-O(DMENQ)_20_ | 512.0 | >512 | >1 | >4* |
| 2C_18_-O(BEDA)_20_ | >512.0 | >512 | >1 | >1* |
| 2C_18_-O(DMENQ)_50_ | >512.0 | >512 | >1 | ≥128 |
| 2C_18_-O(DMEN)_50_ | 512.0* | >512 | >1 | ≥128 |
| 2C_18_-O(BEDA)_50_ | 512.0 | >512 | >1 | >1 |

*Values for this compound showed variability across replicate experiments

**^1^H NMR peak integration analysis for calculation of M_n_ and DP_n_**

The DP_n_ of the DMEN-containing oligomers was calculated using the ratio of an integral associated with protons in the repeat unit to an integral associated with protons in the end-group. The DP_n_ of the 2C_18_-O(DMEN) was determined by using the ratio of the peak “m” at 2.27 ppm (corresponding to the –CH-N-(C**H_3_)_x2_** protons) to peak ‘b’ at 1.26 ppm (corresponding to 56 protons of -C**H_2_** groups in the alkyl tail). This could be summarised using the formula:

𝐷𝑃_𝑛_(2𝐶_18_(𝑂(𝐷𝑀𝐸𝑁)))= $\frac{I_{2.27} \times56}{I_{1.26} \times6}$

The DP_n_ of the BEDA-contained oligomers was calculated using the ratio of an integral associated with protons in the repeat unit to an integral associated with protons in the end-group. The DP_n_ of the 2C_18_-terminated BEDA-containing oligomer was determined by using the ratio of the peak ‘g, h’ from 3.00 to 3.50 ppm (corresponding to the –NH-C**H_2_**-C**H_2_**-NH- protons) to peak ‘b’ at 1.36 ppm. This could be summarised using the formula:

𝐷𝑃_𝑛_(2𝐶_18_(𝑂(𝐵𝐸𝐷𝐴)))= $\frac{I_{3.25} \times56}{I_{1.26} \times4}$

The number average molecular weight (M_n_) was calculated by using the DP_n_ calculated previously for the VDM block, the DP_n_ calculated for the block length of the ring opening reactant and the molecular weight of the initiator.

See below for a representative equation:

𝑀_𝑛_ = 𝐷𝑃_𝑛_ × 𝑀𝑊(𝑉𝐷𝑀)+ 𝐷𝑃 × 𝑀𝑊(𝑅𝑖𝑛𝑔 𝑜𝑝𝑒𝑛𝑖𝑛𝑔 𝑟𝑒𝑎𝑐𝑡𝑎𝑛𝑡)+ 𝑀𝑊(𝐼𝑛𝑖𝑡𝑖𝑎𝑡𝑜𝑟)

The theoretical number average molecular weight (M_n,th_) was calculated by using the DP_n_ calculated previously for the VDM block, the DP_n_ calculated for the block length of the ring opening reactant and the molecular weight of the initiator.

See below for a representative equation:

𝑀_𝑛,𝑡ℎ_ = 𝐶𝑜𝑛𝑣.(%) × $\frac{{[M]}_{0}}{{[I]}_{0}}$ × 𝑀𝑊(𝑉𝐷𝑀) + 𝐷𝑃 × 𝑀𝑊(𝑅𝑖𝑛𝑔 𝑜𝑝𝑒𝑛𝑖𝑛𝑔 𝑟𝑒𝑎𝑐𝑡𝑎𝑛𝑡) + 𝑀𝑊(𝐼𝑛𝑖𝑡𝑖𝑎𝑡𝑜𝑟)
